# Biological Database Mining for LLM-Driven Alzheimer’s Disease Drug Repurposing

**DOI:** 10.1101/2024.12.04.626255

**Authors:** Rico Andre Schmitt, Konstantin Buelau, Leon Martin, Christoph Huettl, Michael Schirner, Leon Stefanovski, Petra Ritter

**Affiliations:** Berlin Institute of Health at Charite - Universitaetsmedizin Berlin, Chariteplatz 1, 10117 Berlin; Berlin Institute of Health at Charite, Universitaetsmedizin Berlin, Chariteplatz 1, 10117 Berlin, Germany

**Keywords:** Alzheimer’s Disease, LLM, Cosine Similarity, AI, Drug Repurposing, Ontology, Large Language Model, Artificial Intelligence

## Abstract

**BACKGROUND:** This study presents a software pipeline that leverages LLMs to apply knowledge stored in natural language (such as in pharmacological texts) and ontologies in a transparent Drug Repurposing (DR) information structure.

**METHODS:** Alzheimer’s Disease (AD) related entries in Gene Ontology and DrugBank were integrated into a Knowledge Graph database to inform LLM prompts. 16,581 drugs were screened for their DR potential by the LLM Llama3:8b. The vector embedding representation of the drugs in the LLM was investigated to asses if LLMs store pharmacological information in alignment with domain expert understanding of pharmacological groups. By measuring the semantic similarity of drugs quantitatively, the performance of the DR pipeline was examined. A manual hallucination check was performed to assess the impact of the ontology-database combination on LLM-hallucination performance. The results were compared against registered clinical trials (RCTs) and proposed medications in meta-analyses to evaluate their predictive value.

**RESULTS:** The embedding analysis showed that the vector representations of drugs in the LLM show clusters in alignment with pharmacological groups. The ontologically enhanced prompt was closer to the expert domain proposals than a zero-shot control prompt without that knowledge. The results of the ontology-based prompt showed fewer hallucinations in their responses compared to the zero-shot control prompting.

**CONCLUSIONS:** Ontology-augmented LLM interaction leads to fewer hallucinations and output closer to expert assessment in comparison with a zero-shot control. We propose retrospective analyses, considering the high-rated drugs and their effect on AD patients as a starting point for further (prospective) research.

## I. Introduction

Over the past decades, data on biological processes and functional relationships in Alzheimer’s disease (AD) and dementia research have proliferated. However, the increased information has not translated into an improved understanding of the disease relative to the degree of the data expansion. In more than 110 years of AD research and over 222,000 studies listed on PubMed on AD, no generally verified AD hypothesis has been confirmed.[1], [2]

While the expanding amount of data on AD and drugs facilitates insights into neural processes[3], it also hinders the synergy of all available knowledge since individual researchers cannot consider the entire knowledge systematically at once. This development requires novel approaches for organizing, accessing, and using data. Perhaps there are applicable drugs for AD treatment that currently have other indications.[4], [5] Such drugs are subject to drug repurposing (DR).[6] With DR, patients may be treated sooner (since repurposed medications do not have to go through a shortened approval procedure),[7] costs for drug development may be reduced, and animal trials can be avoided[8]. Also, the safety profile of drugs that are already in clinical use is more established than the safety profile of an entirely new drug. This paper aims to utilize LLMs in conjunction with ontologies and the DrugBank database to advance data-driven drug discovery. It pays particular attention to the underlying mechanics of how drugs are represented in LLMs and how applying ontologically structured knowledge influences its outputs.

While the DR pipeline is disease-agnostic and could be applied to any other disease, AD is a particularly suitable use case, as there is a wealth of biological data without a clear understanding of the disease and its treatment.

### 1.1 Ontologies and Large Language Models

An ontology is a hierarchically organized representation of knowledge that shows the relation of properties in a subject or area.[9] Domain-specific ontologies have been widely used in biology[10], chemistry[11], and medicine[12] to facilitate the formal organization of data. Ontologies are designed to be interoperable.[9] LLMs may be applied to connect ontologies and knowledge resources.[13] Ontologies and LLMs are complementary technologies. While LLMs are able to process high quantities of data in a way that would otherwise require manual, human work, they may hallucinate and do not work with the newest available data.[14], [15] Hallucinations of LLMs describe output with content that either has no factual basis or is inconsistent.[16] LLMs operate on at least partially outdated knowledge due to their training using world data (usually data on the internet that was available when the model training started). They (often) provide plausible answers by predicting the next most probable token (next word or part of the next word).[17] However, their likelihood of correctness depends heavily on the specific prompt and the knowledge embedded in the model. Scientific data is being published constantly, and it would be challenging to establish an LLM training process that catches up with the scientific progress in real-time. Even in such a setup, hallucinations are still likely to occur the more specific the information one has to ask for. This is due to the setup in which the LLM will always provide some response, as it predicts the most probable next token. Still, the likelihood of a correct or valuable answer becomes lower if the information is a detail that rarely occurs in the training dataset or was not included in the first place.[18]

Those described weaknesses are the fields in which ontologies and other highly structured knowledge resources have strengths and complement LLMs. LLMs may serve as a translation layer. They can transform natural language stored in databases (such as biological processes and pharmacodynamics in the DrugBank) along with semantic knowledge stored in ontologies to standardized output (such as a numeric rating). This rating may then be used in virtual DR screening to advance the research in AD. LLMs represent semantic meaning numerically using embeddings, which are high-dimensional vectors.[19] These embeddings are used to inform the model’s response predictions. As this paper aims to establish an LLM DR pipeline, it must also carefully investigate whether the vector representation of pharmacological terminology accurately reflects plausible domain-specific, pharmacological relationships.

## II. Methods

### 2.1 Methods Overview

As the starting point for the DR pipeline, a database containing information on biological processes (GeneOntology[10] – version 2024-04-24), drugs (DrugBank[20]), and AD pathology, as shown in Figure 1, was created. Gene Ontology (GO) provides a comprehensive collection of biological processes (among other entities).[21] The ARUK-UCL dataset contains GO terms relevant to the development of AD based on the literature, with a focus on microglial proteins and microRNAs involved in AD.[22],[23]

**Figure 1:**
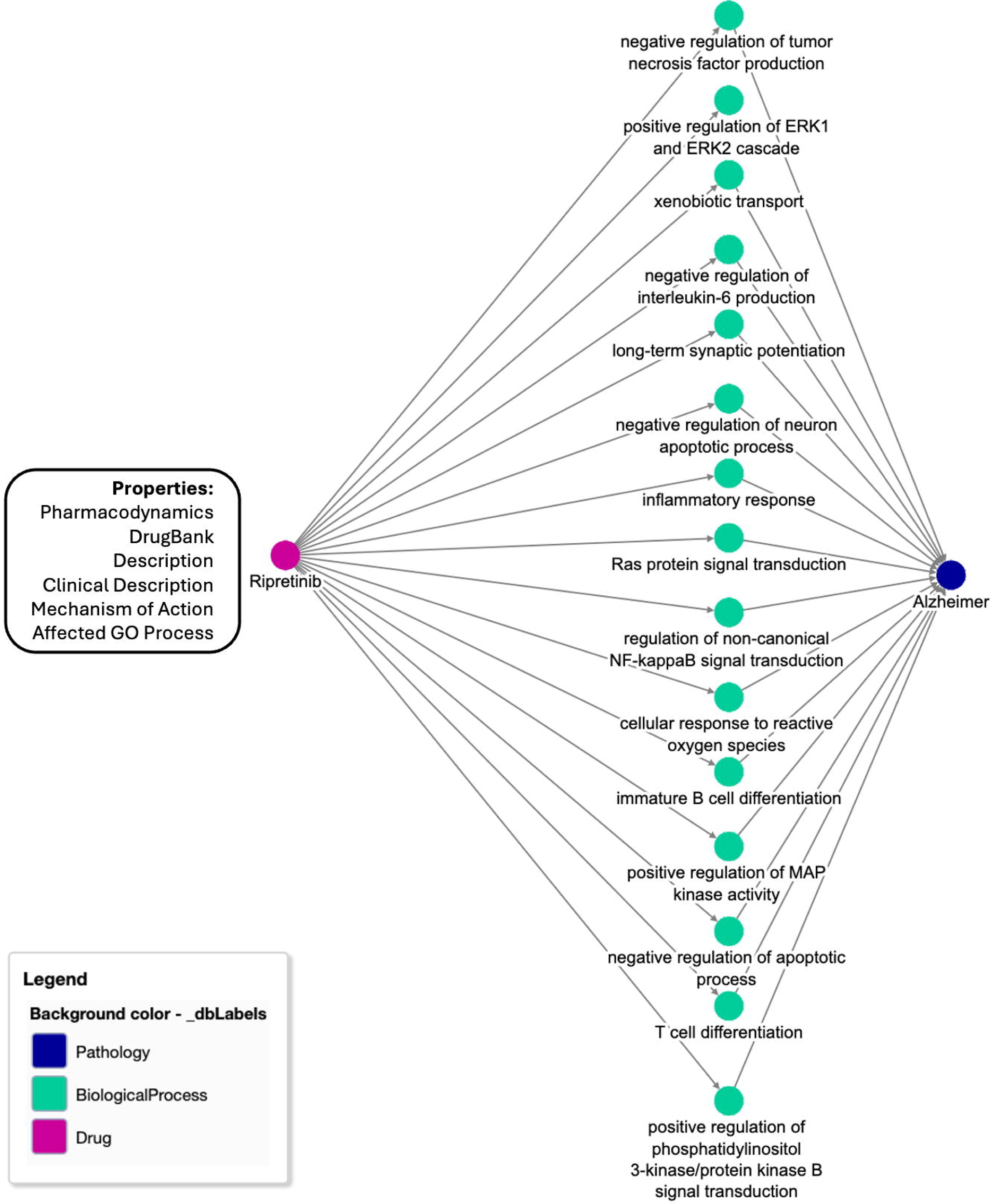
Sample Graph of the Alzheimer’s Drug Repurposing Database This is an example graph from the graph database containing a (manual) selection (15 of 76) of processes (green), AD pathology (blue), and the drug Ripretinib (pink). The arbitrary selection of 15 biological processes was done to enhance clarity and readability in this sample plot (refer to the original code to reproduce the results). The manual selection was necessary for readability on a printed graph, but the full graph can be attained with the code that can be accessed by following instructions *in the open-source GitHub repo (link in ‘Data Availability’ section)*. The graph design can be used to retrieve the Pharmacodynamics, DrugBank Description, Clinical Description (if available), Mechanism of Action, Affected GO Processes, and name of the drug and its associated biological processes (as shown in the figure above). The prompts provided to the LLM followed this principle – a drug and its impacted biological processes are provided along with information about that drug. All the biological processes are associated with AD pathology (according to the ARUK-UCL dataset as described in Methods **1.1**).

Each entry corresponds either to a biological process from GO associated with AD, according to the ARUK-UCL biological process subset, or a drug listed in the DrugBank that has an impact on these processes. This information was embedded in prompts to the LLM to rate the potential of drugs to be repurposed for AD, to get the reasoning for its decisions based on the biological process. That data worked as a translation layer between the natural language-based knowledge in the DrugBank (which contains mechanisms of action, pharmacodynamics, and biological processes described in GO terms) and the formalized biological processes in GO that are connected to the drugs that interact with them. Finally, the drugs with the top 20 ratings of the LLM were reviewed based on the literature published on them to discuss medications that are promising for DR. As control comparison of the ontologically enhanced, structured information prompting a zero-shot control was performed in which the LLM (llama3:8b) only received the prompt asking for a DR rating and the drug. The zero-shot prompt did not contain any information from Drugbank on pharmacodynamics, mechanisms of action, or the biological processes, encoded in GeneOntology terms, that the drug has an impact on, as opposed to the ontologically enhanced prompt that contained this information. To assess the occurrence of hallucinations in both approaches – zero-shot prompt and ontology-enhanced prompt – 50 randomly sampled reasoning responses of the LLM were manually analyzed to identify and count hallucinations/factually wrong outputs of the LLM.

In the next step, we investigated how the LLM, which we used for DR screening, represents the medications we asked about numerically, as an approximation of how semantic relationships were learned by it. As words, such as specific drugs, are represented in an LLM as vectors that can be retrieved by obtaining the embedding of the word, we obtained the embedding representation of the top-performing drugs (the top 50 of the zero-shot control and the top 50 of the ontological prompt).

To comprehend the vectorized semantic relationships visually, we applied the UMAP (Uniform Manifold Approximation and Projection for Dimension Reduction) dimensionality reduction technique. [24] We calculated the cosine similarity between high-performing drugs of each approach to drugs discussed for AD DR in the literature to quantify how close the recommendations of the LLM are to expert assessments.

For biological results interpretation, the drugs were classified by the GO process they impact. To put the ratings into perspective, the ratings of the drugs were compared with those from registered clinical trials on clinicaltrials.gov. This provided insights regarding the relationship between the number of trials and the ratings the drugs received. To test whether the results of the DR pipeline were applicable to all patient groups (ethnicity, nationality, environment) to the same degree, the ratings of drugs were plotted on a map to indicate where the respective drug trials were conducted and what rating the drugs received in each trial. As the LLM was trained on world data (including published drug trial studies), this provided a perspective on how the world population is represented by the studies that provided data for the drugs (world data) considered for DR, and how the LLM responded to the drugs in dependence on where the trials were conducted.

### 2.2 Database Setup

The drugs of the DrugBank and the biological processes (based on the ARUK-UCL dataset) were stored in a graph database. In such a database, information is stored as a network with nodes and their relations to each other, to systematically retrieve information from the graph. Each biological process, drug, and pathology becomes a node connected by its relationships, as shown in figure 1.

The DrugBank contains 16,581 drugs along with extensive information on them, structured by mechanism of action, indication, and other pharmacological key data.(24) There are descriptions of the mechanisms of action of the drugs, as well as the GO processes they impact.(20,25) The indication of each drug is provided as well, which enables the selection of drugs indicated for AD treatment and medications with other indications specifically.

1,778 biological processes from GO and 16,581 drugs from the Drugbank were included. 7,590 drugs in the DrugBank dataset have associated GO-terms and were therefore applicable to be possibly connected with the ARUK-UCL terms. 5,967 drugs were added to the database that had GO terms associated with them that were considered AD-relevant in the ARUK-UCL dataset. All drugs without connections to the (AD-specific) biological processes (10,614) were deleted from the database. Therefore, all drug nodes present in the database at that point had a relationship towards AD-relevant processes, but not necessarily AD as an indication for administering the drug. 44 of these 5,967 drugs either had AD as an indication (such as Memantine(26)) or were under investigation for AD (such as VP025[25], according to the DrugBank dataset. While they were included in the following steps to check their performance in the LLM rating, they are not candidates for DR since AD is already considered to be their (primary) use.

### 2.3 Large Language Model Rating

The LLM was locally hosted for improved accessibility of the data and to be able to reproduce results with the very same model configurations (which may change in a non-open-source cloud-hosted model). Llama3 with 8 billion parameters is an open-source LLM by Meta, suitable for the reproducibility and accessibility of results.[26], [27] The technical details of the LLM are specified in the open-source code (compare ‘Data Availability’ section).

To move towards virtual DR candidate selection, the drug nodes in the graph database were retrieved along with their associated biological process nodes. To get control of model performance without providing the model with structured information (zero-shot prompting for DR), the LLM was simply asked to rate drugs for their DR potential.

For the ontologically enhanced approach of the pipeline, the model was prompted by the properties of the drug node and its biological processes. The LLM worked as a translation layer from the knowledge provided in describing natural language (text in the DrugBank) and standardized knowledge (in GO) to a ranked output. For a drug, its pharmacodynamics, DrugBank description, clinical description, mechanism of action, and name (all-natural language) were provided to the LLM along with the affected GO processes (also natural language but formalized).

The model had to analyze the biological effect of the drug by comparing the GO terms and the data on the drug, and to provide a “reason” why it considers the drug to be a high-potential candidate for DR or not. To reduce the complexity of the dataset and for extended analysis possibilities, the model additionally had to rate the repurposing potential, based on its reason, of each drug on a decimal scale from 0 to 1. The rating and reason were stored in the database. The model was explicitly instructed to base its reasoning and rating on the information it was provided with from the graph database (biological processes impacted by the drug and data on the drug).

All drugs were rated over ten independent iterations of each approach (zero-shot control and knowledge augmented with data from the DrugBank and GeneOntology). Each rating within the ten runs and each run was independent of the other (the LLM did not “know” about the ratings or runs provided before and after). The model did not store any data from any run and was not trained on it either. All iterations were run under equal conditions. The repetitions were performed to account for the variable output of the LLM that provides responses based on probability distributions. Descriptive statistics (minimum, maximum, variance, mean) for each drug across all ten rating iterations were calculated.

We applied the Friedman test for one-way repeated LLM-rating analysis of variance by ranks to test if the ratings per drug were consistent across the rating iterations in each approach (zero-shot prompt control and ontologically enhanced prompt).[28]

Averaging the scores across the 10 iterations led to the final total rating results. Also, it allowed the comparison of each independent run with another to test if the rating was comparable or if it seemed to be random. The authors reviewed the 20 drugs with the highest rating by average for manual review by collecting background information on the drugs in the literature. This included checking if the drugs were already researched for their potential on AD or even indicated for AD (as in both cases, these would not be candidates for DR). The LLM-reasoning output for the remaining drugs was additionally checked for hallucinations (further described in 2.4).

### 2.3 Prompt Engineering

For standardized information processing and drug potential rating of the LLM, we employed chain-of-thought prompting, which has been shown to improve reasoning in LLM decisions.[29] However, the strictly standardized chain-of-thought protocol did not sufficiently discriminate between high and low-potential drugs. Therefore, for the final rating, an associative prompt was used that provided the model with criteria for its rating without forcing it to follow a specific protocol.

We employed a knowledge-augmented approach, utilizing information retrieved from a knowledge graph.[30] This means that the LLM was provided with specific information retrieved from the graph to improve its performance in the ontologically enhanced prompt (such as mechanisms of action and pharmacodynamics of the drug). However, it was provided with that information just once within the prompt, and did not receive additional information while it was generating an output. The zero-shot control prompt, with no knowledge augmentation, did not contain this information.

### 2.4 Hallucination Analysis

To assess the occurrence of hallucinations in both approaches – zero-shot prompt and ontology-enhanced prompt – 50 randomly sampled reasoning responses of the LLM were manually analyzed for each approach to identify and count hallucinations/factually wrong outputs of the LLM.

Hallucination in this context was defined as either:

a. A factually wrong statement in the reasoning AND/OR
b. A self-contradictory statement

Speculative statements (based on correct information from the knowledge-enhanced prompt) were not considered hallucinations, as the task of reviewing biological mechanisms’ impact for possible DR applicability itself is exploratory and, to some degree, necessarily speculative. Results were reported as booleans, representing whether there was either a hallucination concerning the drug during the ratings or not.

In addition to the random sampling hallucination check, the drugs applicable for DR (in the top 20 average ratings of the ontological prompt iterations and no established AD research history), all reasons for the rating that were provided by the LLM were also manually reviewed to identify hallucinations.

### 2.5 Embedding analysis

We assessed whether the vector embedding representation of drugs in the LLM is reasonable by manually curating a list of drugs with different pharmacological groups. We applied the UMAP (Uniform Manifold Approximation and Projection for Dimension Reduction) dimensionality reduction technique.[24] UMAP transformed the 4096-dimensional drug embeddings of Llama to a 2d plot. This allowed us to test whether the pharmacological groups are clustered in a way that aligns with expert knowledge (Figure 4a).

We retrieved the numerical embeddings of the LLM for the top 50 drugs of the zero-shot control approach and the ontologically enhanced prompt approach. The choice of 50 drugs was empirically determined to ensure the clarity and interpretability of the visual pharmacological LLM examination. For a comparison with domain expert knowledge, we employed a list of 22 discussed potential candidates for AD DR in a recent review (Cummings et. al, 2025).[31] The Llama 3:8b knowledge cutoff was in 2023, so it is impossible that it was trained on the review itself and the most recent findings discussed in it.[26]

Taking the top 50 drugs of each prompting approach and the medications discussed in the review provided a plot showing 122 drugs that visualizes how close the drugs of each prompting approach are aligned to the literature comparison (Figure 4b and 4c).

Retrieving the embeddings of the drugs not only allows for visualizing the vector representation of the drugs but also to assess how the zero-shot prompting related to the ontologically enhanced prompt quantitatively. For this exploratory approach, the LLM did not serve as a language production tool but as a semantic representation and mapping technology. By taking the top 50 drugs of the zero-shot control and the top 50 drugs of the ontological prompt, we could calculate the minimal cosine distance between the lists. Cosine distance quantifies the angle between two vectors in a multidimensional space. As words are represented as vectors in LLMs, the cosine similarity can give an insight into how close two terms are.[32] This allowed us (after having checked the plausibility of the approach on the UMAP plot showing the representation of the drugs in the multidimensional vector space) to quantify the distance between zero-shot approach and the ontological prompt to drugs discussed in the literature for AD DR.[31] If we compared the lists binarily we would only get a quantification of either matching or not matching (checking if a drug of the zero-shot control and the ontologically enhanced prompt are discussed in the literature comparison or not).

For example, if on a given list A there was the drug Warfarin (Vitamin K antagonist) and on another list B there was Phenprocoumon (also a Vitamin K antagonist), comparing the lists binarily would mean that there is no match (different drug on each list). However, if Warfarin and Phenprocoumon are represented as vectors, it is possible to quantify the similarity of the drugs under the assumption that the embedding representation and similarity of the drugs are pharmacologically plausible (as tested in the plausibility check described above).

This approach allowed us to measure how close the zero-shot control with the ontologically enhanced prompt is to the drugs discussed in the review. As there is no actual cure for AD yet, we lack a solid ground truth for validating the screening approach. However, by visualizing the top-performing drugs of each prompt on a 2d plot and checking if they are reasonably close to domain expert suggestions, and then calculating their cosine distance, we get a measure of the plausibility of the drug suggestions.

We calculated the minimal cosine distance of each drug on the literature comparison list (meaning the minimal distance we can get for each drug compared to the drugs on the list). The sum of all minimal distances gave a measure of how close the zero-shot control and the ontologically enhanced prompt performed to the literature domain expert suggestions.

Possible distances range from 0 (identical embedding, same drug) to 2 (opposite vector). These possible values originate from the range of cosine similarity -1 to 1.

#### Cosine Similarity

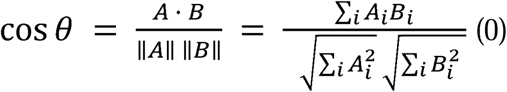

*Cosine similarity range =* [-1, 1] *(1)*

*Cosine similarity = 1 (vectors point in the same direction) (2)*

*Cosine similarity = 0 (vectors are orthogonal) (3)*

*Cosine similarity = -1 (vectors point in opposite direction) (4)*

#### Cosine Distance

*Cosine distance = 1 – cosine similarity (0)*

*Cosine distance = range* [0, 2] *(1)*

*Similarity = 1* → *distance = 0 (identical vectors) (2)*

*Similarity = 0* → *distance 1 (orthogonal vectors) (3)*

*Similarity = -1* → *distance = 2 (opposite vectors) (4)*

These calculations provide a total distance to the literature comparison for each approach (zero-shot control and ontologically enhanced prompt). The total distance is the sum of the minimal distances between each drug of the approach compared to the literature list. The lower the total distance of a tested approach, the closer it is to the literature comparison.

### 2.6 Clinical Trials Data

We complemented the interpretation of the rated drugs by retrieving the registrations of clinical trials from clinicaltrials.gov. This gave the number of studies for each drug and in what clinical trial phase they were registered. In addition, the characteristics of the study population, such as geographic location, were analyzed to address potential bias in the data and LLM rating. The geography of the institutions involved in the studies was then projected on a geographic map to provide insights into the regional distribution of the medications. This step was taken to determine if the results are applicable to all populations to the same degree. Therefore, it was investigated in which regions the drugs were studied in the first place and whether the drug rating is correlated with a geographic region. This approach addressed the biases in the training data of the LLM when advocating for an AI-supported Drug Repurposing approach.

The map represents the number of studies per region and their rating by making use of a color-graded score. The ‘Study Score’ was calculated by the number of studies * rating of drug * 10 to account for the region and study population, and also for the rating the LLM gave for each drug. The *10 in the calculation was done to always get to natural (whole) numbers that are applicable to the plotting.

### 2.7 Reproducibility of Results

The entire study setup was repeated twice to ensure the reproducibility of the results based on the code provided. As described above, the rating of the drugs was repeated ten times to account for randomness in LLM responses and to address the variability of the output statistically (Friedman test). In the replication step, these iterations were only done three times since that process was mainly for setting up the database and the following rating processes. The ten iterations in the final results database themselves are already independent of each other. Therefore, another seven runs in the replication database would not have provided any benefit compared with the actual run of 10 iterations. To further ensure the replication of results, the code is provided openly on GitHub (details in ‘Data Availability’ section). The code is also published and comes along with extensive documentation that allows testing of the results provided in this paper and repurposing for other DR use cases.

## III. Results

### 3.1 LLM Rating Results

The rating distribution between the iterations remained consistent as the Friedman test that tested for differences across rating iterations for each individual drug was insignificant (p-value= 0.15 for the zero-shot control, p-value=0.24 for the ontologically enhanced prompt). This indicates no strong evidence for significant differences in drug ratings across all iterations.

In 135 of 5,967 (∼ 2%) drugs rated by the LLM of the ontological prompt, the input prompt exceeded the context window of Llama (the prompt consisted of more than 8,000 tokens). For 4 (out of 5,967) drugs, the LLM provided at least one non-numeric rating. These four drugs were, therefore, excluded from the Friedman test and the plots based on the rating (but remained in the database).

### 3.2 Best-Performing Drugs

Table 1 lists the 20 drugs with the highest ratings across all iterations in the ontological prompt results. Drugs with AD as an indication scored the highest but are no candidates for DR (as AD is their indication already).

**Table 1:** Top 20 drugs across all iterations with a rating and example reason. This table shows the top 20 drugs of the ontological prompt, along with their average ratings across all 10 rating iterations by the LLM. Also, it provides the variance across the rating and the maximal and minimal rating per drug across the 10 iterations. The reasoning behind the provided rating was taken from the first iteration to give an example. The third column shows the state of research and indication (if applicable) of each drug based on manual literature review. Drugs with no AD-indication or AD research background are indicated by purple background as they are potential drug repurposing candidates. The decision to set ‘20’ as the upper limit for drugs being analyzed more closely was done arbitrarily but based on the the distribution of ratings. Drugs applicable for DR (as they are not under investigation for AD) are marked in purple.

| Name | Average Rating | Variance | Max | Min | Reasoning in the first iteration | State of research background |
| --- | --- | --- | --- | --- | --- | --- |
| **CTS-21166** | 0.90 | 0.00024 | 0.95 | 0.9 | CTS-21166, as a beta-secretase inhibitor, targets the proteolytic process involved in Alzheimer's pathology. The biological processes impacted by CTS-21166, such as protein catabolic process and membrane protein ectodomain proteolysis, are relevant to driving Alzheimer's disease. Additionally, the directionality of the impact suggests a preventive or curative effect on Alzheimer's. With excellent pharmacokinetic properties, it is highly likely that patients can take this drug to prevent or treat Alzheimer's. | Under investigation |
| **Lecanemab** | 0.90 | 0.00191 | 0.95 | 0.8 | Based on the information provided, Lecanemab targets amyloid-β plaques in a dose- and time-dependent manner, which is highly relevant for driving Alzheimer's disease. The directionality of its impact suggests a preventing or curing effect on Alzheimer's. Additionally, it can be taken by patients to prevent or treat the disease, as demonstrated by its clinical trials and FDA approvals. | AD |
| **Verubecestat** | 0.88 | 0.00224 | 0.95 | 0.8 | Verubecestat's mechanism of action in inhibiting BACE, a key enzyme in amyloid $\beta$ peptide production, suggests a potential therapeutic effect on Alzheimer's disease. The biological processes it impacts, such as proteolysis, protein catabolic process, and membrane protein ectodomain proteolysis, are relevant to driving Alzheimer's pathology. | Was investigated |
| CAD106 | 0.84 | 0.00266 | 0.9 | 0.8 | The drug CAD106 has a mechanism of action that directly targets the formation of beta-amyloid plaques in the brain, which is considered a hallmark of Alzheimer's disease. This suggests a strong potential for treating the underlying pathology. The impacted biological processes are also relevant to Alzheimer's disease, including neuron remodeling and apoptosis, which supports the drug's potential for preventing or curing the disease. | Under inve |
| NGX267 | 0.84 | 0.00266 | 0.9 | 0.8 | NGX267 has a high potential for repurposing as it targets biological processes relevant for driving Alzheimer's, such as synaptic transmission and signal transduction. Additionally, its mechanism of action on phospholipase C-activating G-protein coupled receptor signaling pathway may contribute to delaying disease progression. Furthermore, NGX267 has shown potential to reduce symptoms and slow disease progression in Alzheimer's patients. | Was inv |
| 4-{2-[(7-amino-2-furan-2-yl[1,2,4]triazolo[1,5-a][1,3,5]triazin-5-yl)amino]ethyl}phenol | 0.83 | 0.00233 | 0.9 | 0.8 | The drug has a positive impact on biological processes associated with Alzheimer's disease, including regulation of synaptic transmission and plasticity, which are key mechanisms underlying the progression of Alzheimer's. Additionally, it affects apoptosis and inflammatory responses, both of which are important in Alzheimer's pathology. However, some of these processes are not specific to Alzheimer's, such as membrane depolarization and locomotory behavior. Nevertheless, the overall profile suggests a potential therapeutic effect on Alzheimer's disease. | Re<br>uni |
| Brasofensine | 0.83 | 0.00233 | 0.9 | 0.8 | Brasofensine's ability to positively regulate ERK1 and ERK2 cascade, long-term synaptic potentiation, neuroblast proliferation, and receptor internalization suggests its potential to promote neuronal health and prevent Alzheimer's disease pathology. | Was inv<br>Parkinson' |
| Ripretinib | 0.83 | 0.00788 | 0.95 | 0.7 | The drug's ability to positively regulate peptidyl-serine phosphorylation, which is relevant to Alzheimer's disease, and its impact on cellular response to thyroid hormone stimulus, which may be related to the disease's underlying biology. | Treatment<br>gastrointes |
| CX-717 | 0.83 | 0.00233 | 0.9 | 0.8 | CX-717's impact on glutamate receptor signaling pathway, synaptic transmission, and signal transduction suggests potential therapeutic benefits for Alzheimer's disease treatment. | Under inve |
| D-glucose | 0.82 | 0.00736 | 0.95 | 0.7 | The drug D-glucose is a glucose transporter that regulates glucose transmembrane transport, which is involved in the insulin receptor signaling pathway and has been linked to Alzheimer's disease. Additionally, it regulates calcium ion import, which can have neuroprotective effects. | investiga<br>considered |
| Aranidipine | 0.82 | 0.00402 | 0.9 | 0.7 | The drug Aranidipine targets biological processes related to receptor-mediated endocytosis, retina homeostasis, and transmembrane transport, which are implicated in Alzheimer's disease. Furthermore, it modulates inflammatory response, negative regulation of tumor necrosis factor production, and negative regulation of interleukin-6 production, all of which are key features of the disease. Given its potential to influence multiple relevant biological processes, I believe Aranidipine has a high likelihood of being effective in Alzheimer's disease. | Under inve<br>hyp |
| **Pertuzumab** | 0.82 | 0.00180 | 0.9 | 0.8 | Pertuzumab's mechanism of action involves the inhibition of HER2-mediated signaling, which is also implicated in Alzheimer's disease. Additionally, Pertuzumab has been shown to have neuroprotective effects and may help prevent neuronal degeneration. The biological processes it impacts, such as signal transduction, protein phosphorylation, and positive regulation of cell growth, are all relevant to the pathology of Alzheimer's disease. | Indicate positive meta |
| **JNJ-37822681** | 0.82 | 0.00180 | 0.9 | 0.8 | JNJ-37822681's ability to positively regulate ERK2 cascade, long-term synaptic potentiation, and neuroblast proliferation suggests its potential to modulate Alzheimer's disease-related pathways, warranting further testing. | Inv<br>S<br>tr |
| **Umifenovir** | 0.82 | 0.00236 | 0.95 | 0.8 | Umifenovir regulates lipid metabolic process, which is associated with Alzheimer's disease pathology, making it a potential candidate for repurposing. | Antiviral age<br>in some<br>pro<br>treatment<br>and other res<br>in |
| **CYT006-AngQb** | 0.82 | 0.00124 | 0.9 | 0.8 | The drug CYT006-AngQb has a significant impact on biological processes related to brain development, cell surface receptor signaling pathway, and intracellular signal transduction, which are all relevant to Alzheimer's disease. Additionally, it regulates inflammatory response, which is also a key feature of the disease. Furthermore, it positively regulates transcription, DNA-templated, phosphoprotein phosphatase activity, nitric oxide biosynthetic process, extrinsic apoptotic signaling pathway, cytokine production, and protein metabolic process, which are all potential mechanisms that could be targeted to treat Alzheimer's. | A therapeu<br>develop<br>hyp<br>activating<br>syst<br>Angi |
| **DPI59** | 0.82 | 0.00344 | 0.9 | 0.7 | DPI59's ability to regulate cell cycle, protein autophosphorylation and Ras protein signal transduction makes it a promising candidate for Alzheimer's disease kinase B signaling repurposing. | Re |
| **MK-0354** | 0.82 | 0.00177 | 0.9 | 0.8 | MK-0354's ability to positively regulate release of sequestered calcium ion into cytosol, protein phosphorylation, and phosphatidylinositol 3-kinase/protein kinase B signal transduction, which are all relevant to Alzheimer's disease regulation of epidermal growth factor receptor signaling pathway. | Was inv<br>Athero |
| **Indigotindisulfonic acid** | 0.82 | 0.01566 | 0.95 | 0.5 | The drug Indigotindisulfonic acid has shown promise in regulating neuron differentiation and MAPK cascade, both of which are relevant biological processes for Alzheimer's disease. Additionally, it modulates G-protein coupled receptor signaling pathway, a key pathway involved in neuronal function and survival. Given its potential to impact multiple key pathways, I believe this drug is worth further testing for repurposing as an Alzheimer's treatment. | Contra<br>cardiovascul |
| Darifenacin | 0.82 | 0.00177 | 0.9 | 0.8 | Darifenacin's ability to modulate the cholinergic system, which is involved in Alzheimer's disease pathology, makes it a strong candidate for repurposing. Additionally, its impact on lipid metabolic process and xenobiotic transport may also contribute to its potential therapeutic effects. | M3 muscarinic treatment |
| Selenious acid | 0.82 | 0.00177 | 0.9 | 0.8 | The drug Selenious acid has a high potential for Alzheimer's disease drug repurposing due to its impact on biological processes related to apoptosis regulation, inflammation modulation, and oxidative stress response. Its ability to regulate neuron apoptotic process, negative regulation of release of cytochrome c from mitochondria, and negative regulation of oxidative stress-induced intrinsic apoptotic signaling pathway suggests that it may have a therapeutic effect in reducing neuronal death and oxidative damage associated with Alzheimer's disease. | Essential treatment under investigation for AD treatment |

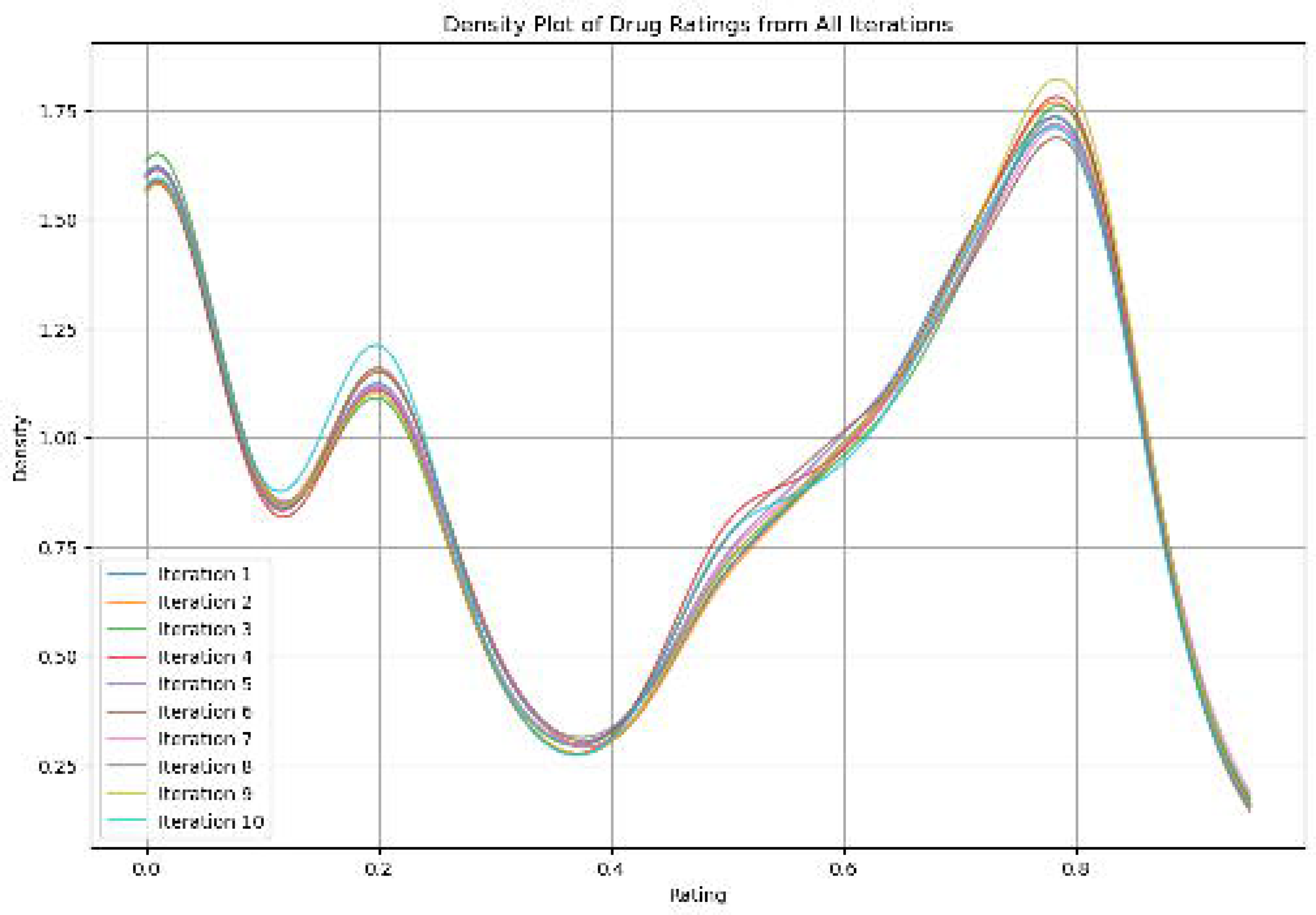
2a) This graph compares the distribution of drug ratings across 10 iterations per individual drug. ‘Iteration 1’ to ‘Iteration 10’ Each rating was independent of the others, and the LLM was not aware of its previous rating. This comparison was made to check for randomness of the LLM rating of each drug. In most cases, the rating provided by the LLM was visually similar for each iteration as it can be seen on the density plot. This visual impression is in agreement with the Friedman-Test that indicated no significant rating differences between the iterations for each individual drug. 2b) This graph shows the distribution of each individual drug’s rating sorted by their average ratings. Each drug has a boxplot assigned, that shows the distribution of ratings across the 10 rating iterations. The x-axis shows the average rating of each drug. The opacity of each boxplot depends on the range of values (larger boxplots are more transparent) to allow differentiation between the boxplots as those drugs with small interquartile ranges would otherwise be covered. This graph has to be considered along with the Friedman test that tested for significant rating differences of e drug over all rating iterations that did not suggest a significant difference.

CTS-2166 is a beta-secretase inhibitor under investigation for AD disease treatment.[34] Lecanemab is an amyloid beta-targeting antibody approved by the FDA for AD treatment.[35] Verubecestat was under investigation for AD treatment[36]; its development, however, was put on hold in 2017.[36] The drug with the highest rating that is not under explicit investigation for AD disease is Ripretinib, a tyrosine kinase inhibitor indicated for the treatment of advanced gastrointestinal stromal tumors.(41) Ten drugs of the 20 highest rating medications have neither been indicated for AD nor been under investigation for it.

Nine of the 20 best-performing drugs have either been under investigation or were investigated for Alzheimer’s disease. In two cases (‘4-{2-[(7-amino-2-furan-2-yl[1,2,4]triazolo[1,5-a][1,3,5]triazin-5-yl)amino]ethyl}phenol’ and ‘DPI59’, the state of research was not determinable based on the literature. Nine high-performing drugs had no determinable history of AD consideration.

**Figure 3:**
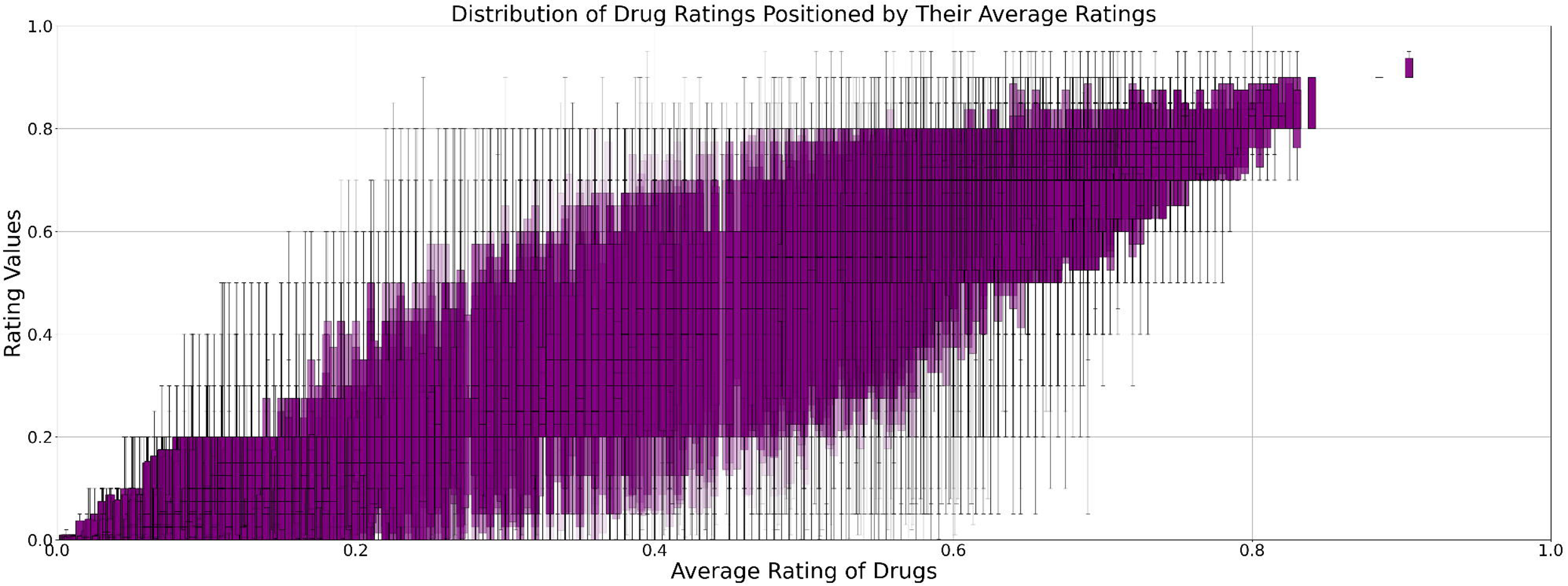
Distribution of Top 20 Drug Ratings Across Rating Iterations This plot shows the boxplots of the upper 20 drugs sorted by their average rating. As some of the drugs had the same average rating (such as Lecanemab and CTS-21166) the sorting for those drugs is random. Indigotindisulfonic acid has the highest variance at 0.01566. The drug shortened for layout reasons in the plot is: ‘4-{2-[(7-amino-2-furan-2-yl[1,2,4]triazolo[1,5-a][1,3,5]triazin-5-yl)amino]ethyl}phenol’.

### 3.3 Hallucination Check

The random sampling of 50 reasonings for the zero-shot control and the ontological, knowledge-enhanced prompt each provided 100 LLM reasoning to be checked manually for hallucinations. 19 out of 50 zero-shot control responses contained hallucinations, whereas there was only one hallucination in the ontology-based prompt. In addition to the random sampling, the drugs highly rated in the ontology-based prompt were checked for hallucinations.

One hallucination was detected across all rating iterations of the DR candidates (Pertuzumab). The LLM reported, “Pertuzumab has been shown to have neuroprotective effects and may help prevent neuronal degeneration,” which is a wrong statement.[57]

While D-Glucose does not fall in the category of DR-applicable drugs as it has an AD research history and is not considered a treatment,[46] it has to be pointed out that a hallucination occurred for D-Glucose as well (provided in the reasoning in Table 1). The LLM called D-Glucose a “glucose transporter,” which is factually wrong.

### 3.4 Embedding analysis

The assessment of the pharmacological plausibility of the vector representation in the LLM embeddings was first performed with a manually curated list of drugs belonging to shared pharmacological groups. While the embeddings are in 4096 dimensions, the UMAP dimension reduction technique allowed us to visualize how the drugs are represented in vector space on a 2d plot (Figure 4a). The drugs are mostly clustered by their pharmacological group, which makes the vector representation pharmacologically reasonable. The closer relationship of the ontologically enhanced embeddings can also be visually seen on the UMAP dot visualization that reduces the high dimensionality (4096 dimensions of Llama3:8b) to a 2d plot (Figure 4b and 4c). Visually, it is apparent that the drugs with a high rating in the ontology-based prompting (orange in 4b and 4c) are closer to the literature comparison (green) than the top 50 drugs of the zero-shot control (blue).

**Figure 4.**
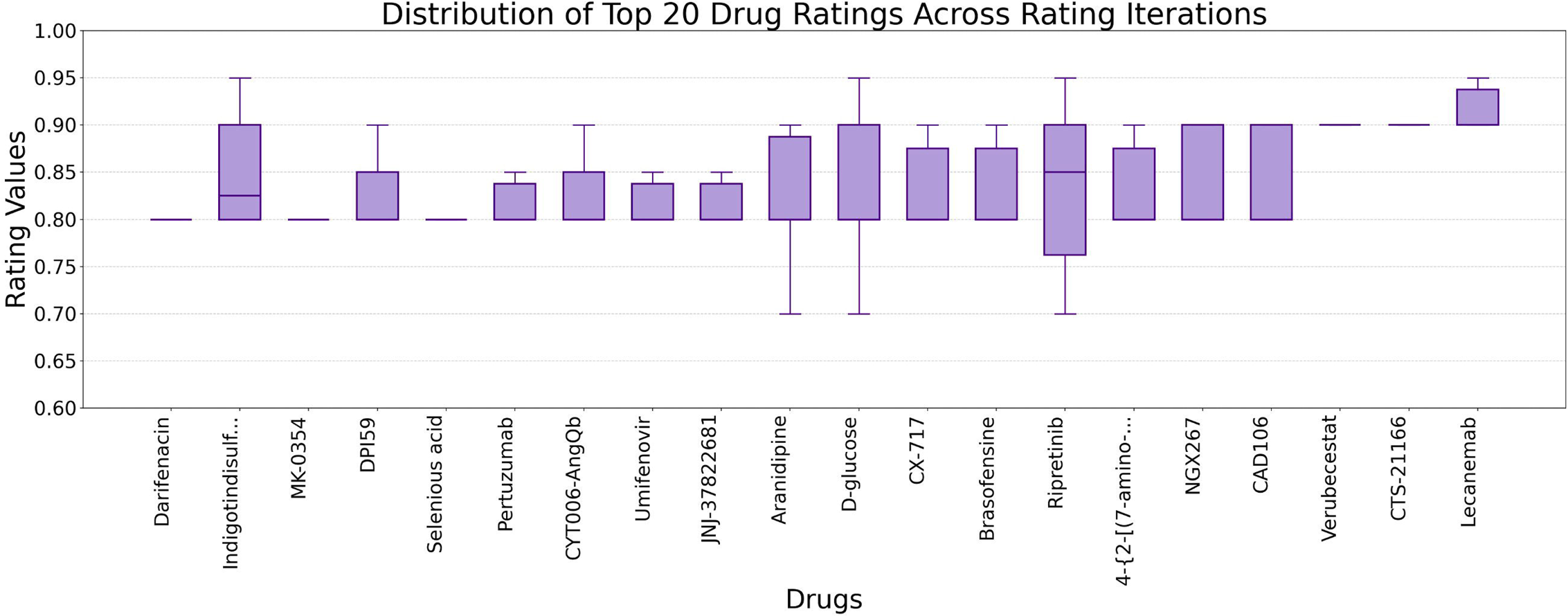
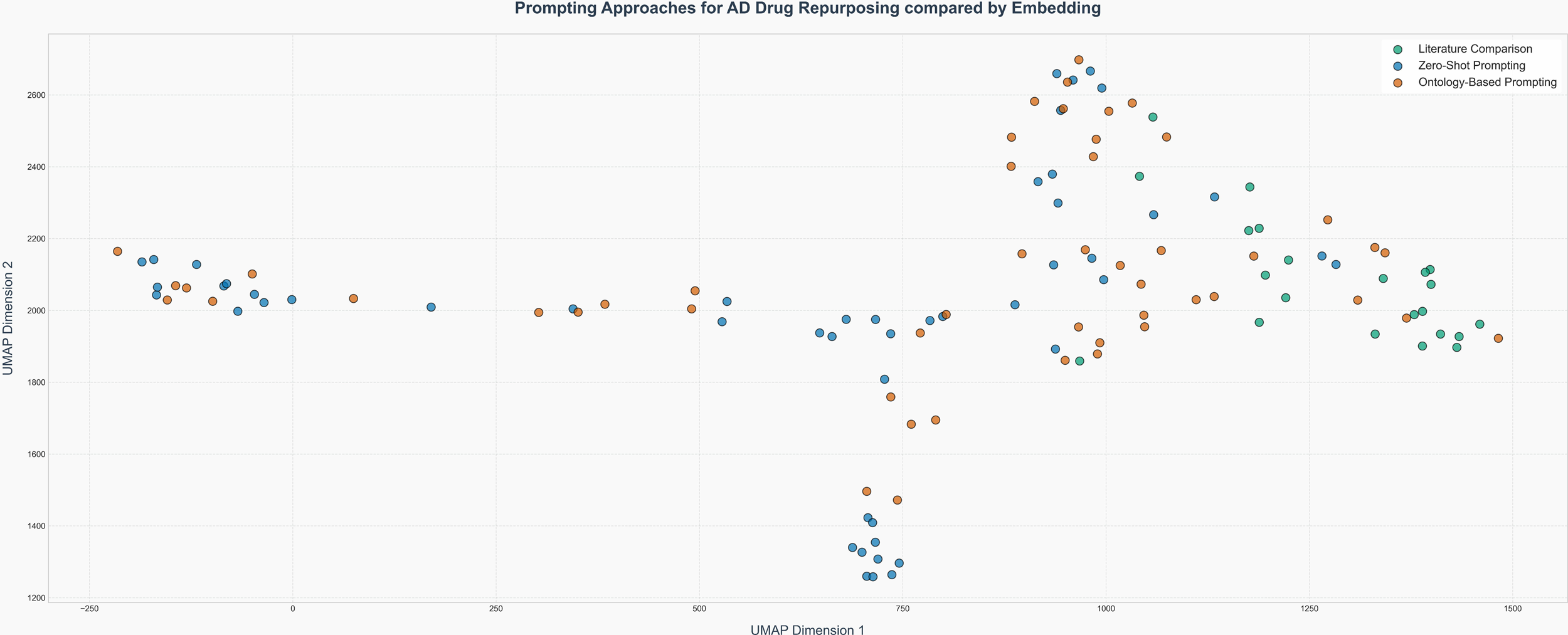
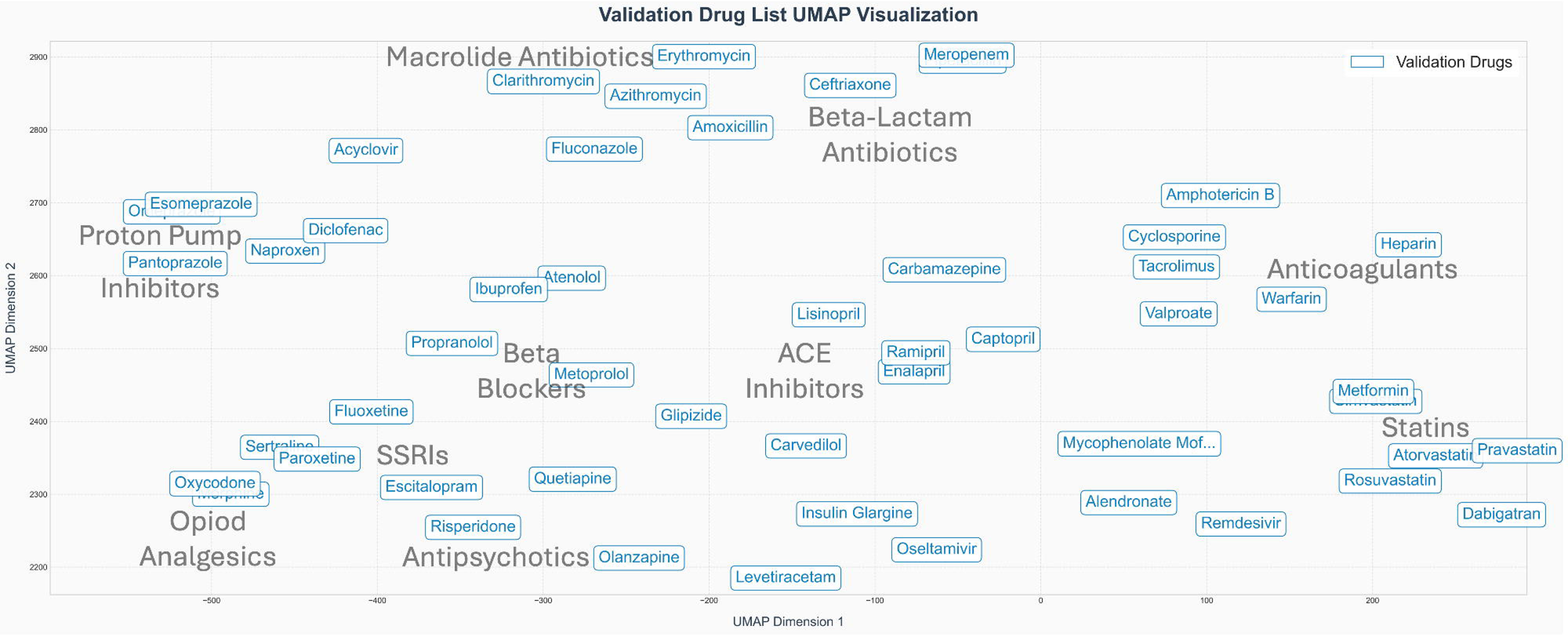
a) UMAP vector representation of a manually curated list of drugs belonging to pharmacological groups to assess the pharmacological plausibility of embedding storage of drugs. The UMAP technique reduced the embeddings of the drugs from 4096 dimensions to a 2d plot. Drugs that belong to the same pharmacological group are also close to each other in the vector space. Figure 4 b). Vector representation of highly rated drugs by each prompting approach compared with drugs proposed in the literature, plotted on a 2d plot through UMAP dimensionality reduction without drug names. This plot shows the embeddings of the top 50 drugs of each approach as a scatterplot. The drug names are not shown on this plot for visual clarity. Figure 4 c). Vector representation of highly rated drugs by each prompting approach compared with drugs proposed in the literature, plotted on a 2d plot through UMAP dimensionality reduction with drug names. These are the embeddings of he top 50 drugs of each approach projected with the actual names on the dimensionality reduced plot. This shows that drugs of similar pharmacodynamics are also clustered together in the high-dimensional vector embeddings of the LLM. Drugs that do not have a generic name but only an IUPAC classification (long chemical names) are also clustered.

The cosine similarity between the top 50 drugs of the zero-shot prompting and the ontology-enhanced prompt to the literature comparison was calculated. The average distance of the ontological prompt was closer (0.295) to the literature comparison than the zero-shot prompt (0.376), as shown in Table 2.

**Table 2:**
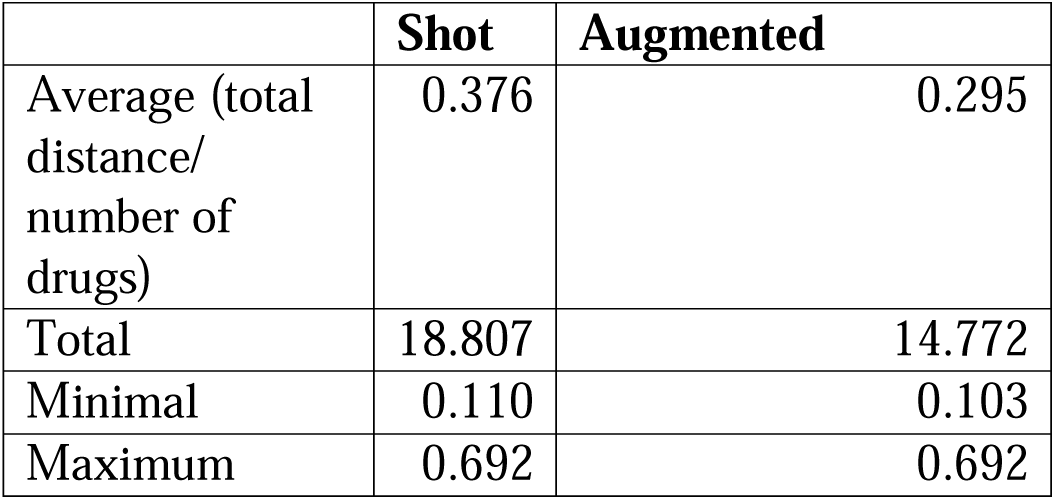
Cosine distances between Zero-Shot control prompting and Ontology Augmented Prompting.

### 3.5 Studies on the Drugs

We retrieved data for all drugs associated with AD from clinicaltrials.gov (on 07/26/2024). However, of the 5,967 drugs with AD-associated GO processes, only 2,918 had retrievable trials available. The data contained information on the phase of the drug trial and the geographic location of the research facilities investigating the drug (such as a hospital).

**Figure 5:**
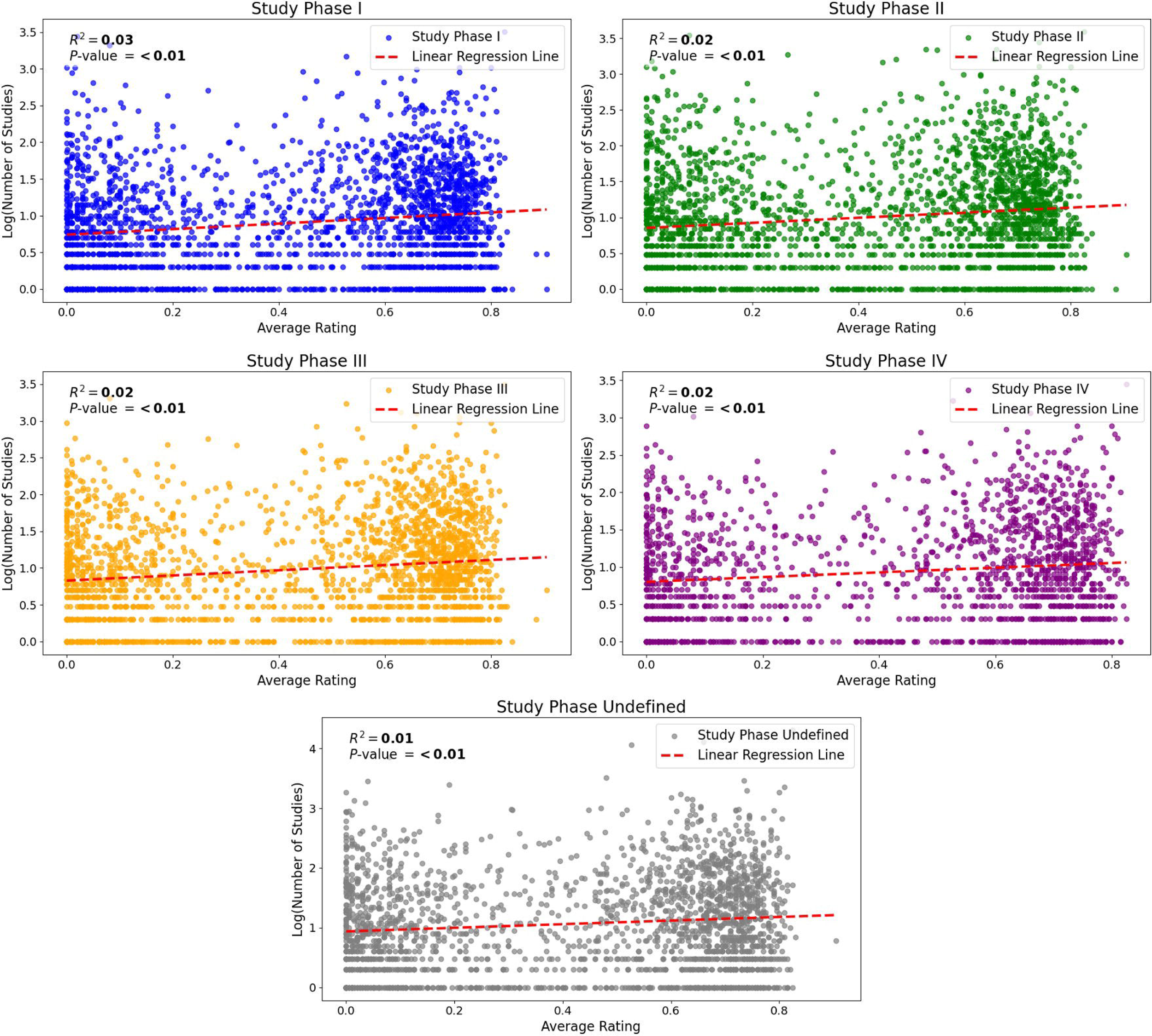
Rating of Drugs in Dependence of Study Phases and Number of Studies. Each plot shows the base-10 logarithm of the number of studies associated with each drug in dependence on the average rating the drug received across all rating iterations. Each subplot stands for one Study Phase according to the registration of the studies on clinicaltrials.gov. A linear regression line was fitted to the individual data points using the least square method. The analysis suggests a tendency towards higher average ratings for drugs that have a greater (log-transformed) number of (registered) studies associated with them. For all phases, the p-values associated with the regression coefficients were lower than 0.05, indicating that the relationships are statistically significant at the 5% significance level. However, the R² values for all regression models were also lower than 0.05 showing that less than 5% of the variance in the log-transformed number of studies is explained by the average drug ratings. Consequently, while the linear regression suggests a significant relationship, the average drug rating is a weak predictor of the number of studies associated with each drug. As there were only 2918 (retrievable) registered studies for the drugs investigated in this study, 3049 drugs are not shown on the graphs.

## IV. Discussion and Conclusion

### 4.1 Summary of Main Results

A systematic network creation was performed that unifies information from biology (GO), pharmacology (DrugBank), and their interplay. 5,967 drugs were rated for their DR potential. The rating of the individual drugs did not vary significantly (according to the Friedman test conducted) across the rating iterations of the LLM (both in the zero-shot control prompting and the ontologically enhanced prompt). The ratings remained fairly stable across all iterations. The highest-rated drugs of the ontological prompt were either indicated or investigated for AD treatment. However, in the upper twenty drugs, ten drugs were applicable for DR consideration since they have not been associated with AD in the literature.

The embedding analysis of the highly rated drugs gave an insight into how the LLM stores the drugs numerically. Although we visualized this representation that is stored in 4096 dimensions by Llama 3:8b, it was possible to see pharmacologically meaningful clustering of the drugs (e.g., monoclonal antibodies and GABAergic medications grouped as closer to each other than unrelated mechanisms of action, as visible on plots 4a, 4b, and 4c.

While the validation plot (4a) strongly suggests that the closeness in the embedding space is a proxy of comprehension (as far as one can speak of comprehension in an LLM) of the pharmacological groups, other factors probably also play a role. Phonetic similarity and shared suffixes belong to the common nomenclature in pharmacology. The fact that the monoclonal antibodies in Figure 4c are grouped close together may be related to the LLM being trained to associate this pharmacological group close to each other. However, it is indeterminable to differentiate how much the shared suffix “mab” has contributed to such clustering. However, in the validation plot 4a, we also saw Fluoxetine and Escitalopram (both SSRIs) grouped together, which indicates that the embeddings do not rely solely on shared phonetics. Besides phonetic similarity, long medication names that the LLM has possibly never encountered during training seem to have caused a cluster in the highly rated drugs, too (Figure 4c). The cluster of drugs that have no generic name but an IUPAC classification only, although they are chemically very different, shows that the LLM categorizes these drugs similarly to how a human would read them (recognizing that this is a 20+ word long IUPAC classification instead of classifying the pharmacological group itself). However, some of these drugs got consistently high ratings by the LLM (both in the zero-shot prompting without background information and the ontological prompting that entailed such data). From these results, it is indeterminable if these drugs were highly rated because the LLM had no appropriate response tokens, stochastically, as there are many such IUPAC classified drugs in the Drugbank, and the likelihood that some of them would be highly rated was high, or if these drugs are actually highly promising AD DR candidates.

The results from the cosine distance calculation are in alignment with the visual representation of the highly rated drugs. The ontology-based, knowledge-augmented approach is closer to the drugs applicable for AD DR that are discussed in the literature. Also, hallucinations occurred fewer (1/50) when the LLM was prompted with the ontology and database retrieved knowledge compared to the zero-shot control (19/50).

The number of registered drug trials had a slightly increasing impact on the rating of the LLM, which may be caused by more training data on these drugs. Drugs with high ratings and many associated trials were mostly investigated in North America and Europe, which may indicate that the results are not applicable to all patient groups equally.

While these results need to be treated with caution, as we have no ground truth to validate them, they strongly suggest that an ontology-supported, knowledge augmented prompt performs more similarly to an expert evaluation than the zero-shot control.

### 4.2 Drug Research Facilitation

Due to the graph database design, results can be queried quickly and deployed for individual project purposes. This may include domain-related queries for specific use cases or an improved understanding of the interplay of drugs in polypharmacy if different medications target the same biological processes.

While the current results do not justify prospective DR clinical trials, we propose conducting retrospective analyses to evaluate the performance of high-rated drugs and their impact on AD patients. Ripretinib (ranked 8th) was the highest-rated drug with a determinable research history not investigated for AD. Ripretinib is a tyrosine-kinase inhibitor. Tyrosine-kinase inhibitors were recently proposed as candidates for AD DR in the literature.[4]

Analyzing the effect of Ripretinib and Pertuzumab – the highest-rated medicines that are not AD-related by their indication - on patients across AD stages can be a starting point for this. However, limiting such a study to those two drugs is arbitrary, and additional high-rating drugs may be considered, too. If a retrospective study indicates a significant, positive effect on AD for some of the high-rated drugs, those medications may be tested in RCTs. Actual drug trials that test the proposed medications are the only way to validate the concept.

The LLM-driven DR pipeline may also be employed for other pathologies as long as there is a systematic biological entity set for accessing biological data. This only requires a set of biological processes from GO that can then be matched with drugs that have an impact on them. This set of biological processes can either be manually curated by experts or a given set can be applied.

While we do not recommend prospective studies based on these exploratory results, if clinical trials are considered in the future, a thorough safety profile check has to be done.

### 4.3 Limitations

Since the AD driving processes are not yet known/fully understood, the ARUK-UCL dataset, which was used as a starting point, may lack important GO terms or also include terms that are not significant to the pathology’s etiology. While the DrugBank is one of the most established datasets for drugs, it may not include all experimental and new drugs that are possibly applicable for drug repurposing for AD. Also, only 7,590 of the drugs have annotated GO terms. Those without annotated GO terms were excluded from our approach since it was based on the applicability of the preselected biological terms from ARUK-UCL. While we tested different prompts to find one that brings valuable outputs from the LLM, prompt engineering is always imperfect since the number of possible prompts is infinite. ‘*rating_JSON_generator.py’* in the GitHub repo shows how the prompts were generated (see ‘Data Availability’ section for details).

The Large Language Model we used (Llama3:8b) had 8 billion parameters and a context length of 8.2k tokens. Such a model would have been considered highly advanced only a few years ago, but larger models have emerged. In the future, LLM-driven drug repurposing should also consider such models. However, one also has to keep in mind that a larger model is not equivalent to better outcome performance, and improved results based on a larger model are not guaranteed at all.[60] As we have seen in the embedding analysis, smaller models may not be able to represent specific drugs in their vector space (e.g. drugs with IUPAC classification only). Currently, the number of openly available LLMs is limited, and this study only used Llama3 by Meta and no models by other providers. Future research should extend the model selection to a diverse set of open-source LLMs.

The LLM-driven DR approach itself is debatable since addressing DR with an LLM is not established. LLMs work by predicting the next likely token (usually a word or a part of a word). They have no consciousness of the matter they are dealing with and are easily impacted by keywords and minor variations in the prompt that is addressing them. Further research that analyses the scientific value of the ratings is required to validate such an approach.

The upper seven drugs that had the highest rating in the structured input approach for DR potential were either indicated for AD already or were/are at least under investigation for treating AD. Since the LLM was prompted to rate all drugs that had AD-driving biological processes associated with them, some of the drugs were necessarily under investigation for AD or even indicated for it in some countries (such as Lecanemab). Also, in cases where the indication was provided in the DrugBank dataset, the LLM received that information as part of the prompt.

The fact that AD-associated medications had the highest rating allows for several interpretations. While it may indicate that the LLM is aligned with pharmacological AD research, it may also have rated these medications so highly because it was trained based on world data (which likely contained information regarding these medications). Whatever the specific reason may be why these drugs received the highest ratings, it would have been unexpected if they were, as AD drugs, rated as being less promising than (currently) unrelated drugs. Future LLM DR research using more advanced models that can handle more input than current models may benefit from providing more details about each drug.

The comparison of the LLM approach outputs with the literature data of a review was performed due to the lack of a ground truth. The very point of DR is to discover causalities that are not yet understood, so when trying to establish an LLM DR pipeline, it has to be part of the research to assess suggestion quality while lacking a ground truth to compare to. We addressed this by matching different lists semantically (using the vector embeddings of the high-performing drugs) to test if the ontology-based prompting, which provided the LLM with information it would not have had otherwise, brought the LLM output closer to domain expert assessments. This approach is exploratory, and cosine distance may not be the most suitable measure to assess similarity.[32]

The drug classification based on the GO term for Figure 6 was based on the classification of the LLM since there were no pharmacological class categorizations available for many medications (particularly those that are still so experimental that such data is not available). Mistakes in terms of classification may have occurred here. However, since the figure was about giving an overview of the drugs with their rating, a few single wrong classifications would – given the scale - have changed neither the results nor the graph notably.

**Figure 6:**
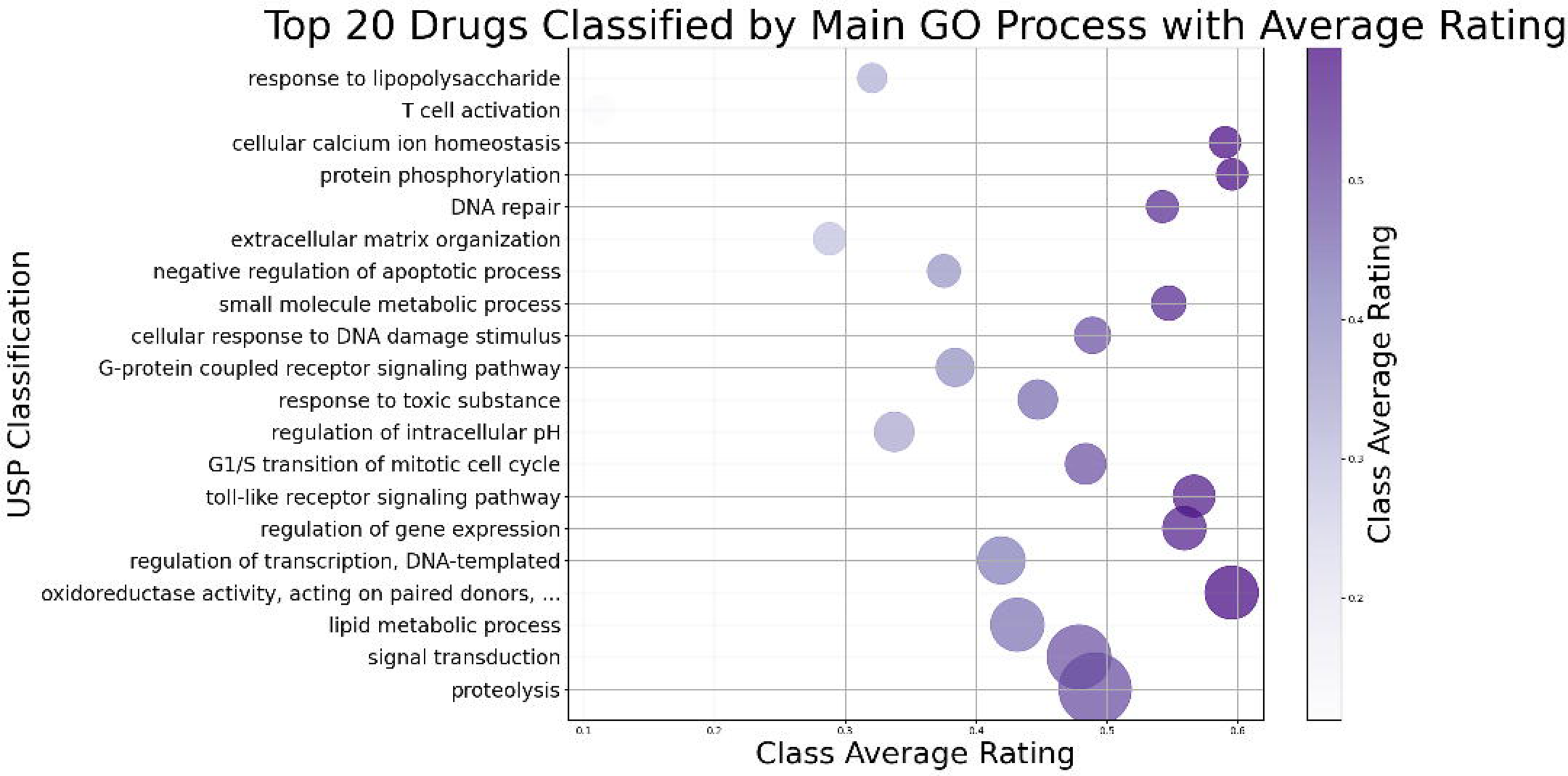
Classification of Rated Drugs by Gene Ontology (GO) Process. In this plot, the drugs were classified by the LLM according to the GO process that was most relevant for the drug repurposing (DR) (based on the reason provided). The size of the bubble depends on the number of drugs whose AD treating mechanism was considered mostly driven by the respective GO process. The x-axis shows the average rating of drugs classified with that GO term. The average rating is also color-graded, deeper purple indicating a higher class average rating. Only the 20 most classified GO-terms are shown for readability purposes. The entire process on how the graph was attained is described in the original code (consult Appendix A for replication) and on the online documentation. The full GO-term at the fourth position from the bottom (which was truncated for layout reasons) is: ’oxidoreductase activity, acting on paired donors, with incorporation or reduction of molecular oxygen, reduced flavin or flavoprotein as one donor, and incorporation of one atom of oxygen’. The geographic data concerning the rating of the LLM led to a map that has most of its data points in North America and Europe. Each dot represents a research-facilitating institution (such as a hospital). Most retrieved studies were conducted on those continents, but the LLM rating for drugs tested in these areas was also higher. The color density depended, as described in the Methods, on the number of studies per area and the average rating of the drugs.

**Figure 7:**
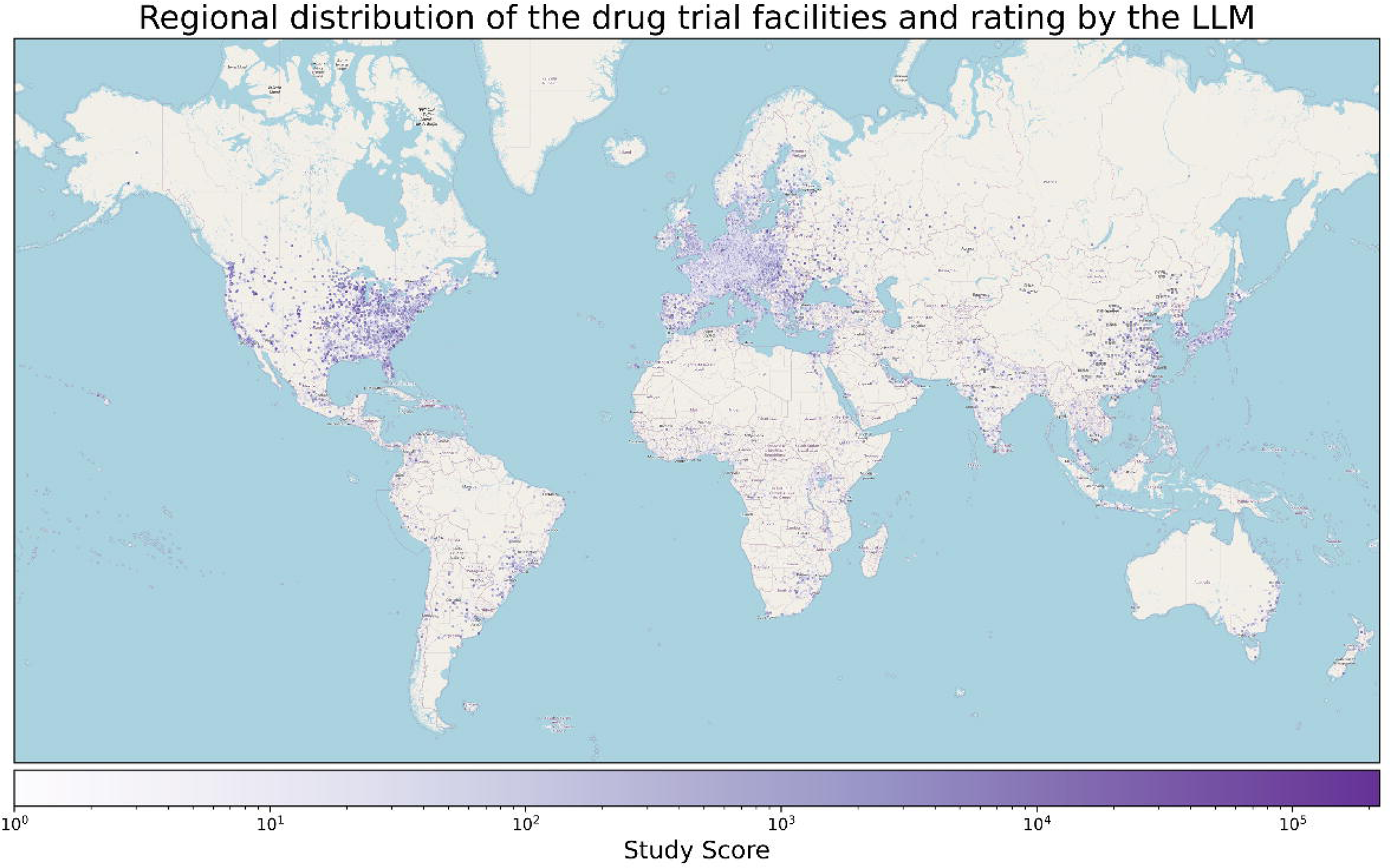
Regional distribution of the drug trial facilities and rating by the LLM The map[58], [59] shows the regional distribution of the drug trial facilities and the LLM’s ratings. Each dot on the map represents a drug trial facility provided in the trial registrations. The color intensity corresponds to the ‘Study Score’, which was calculated using the number of studies * average rating * 10 to obtain natural (whole) numbers. Therefore, darker purple indicates higher ratings and/or more conducted studies in an area. For multicenter studies (e.g., several hospitals), all research facilities are shown on the map. A logarithmic normalization was applied to the ‘Study Score’ to allow for color differentiation. This provides a more interpretable representation of the score across all regions. The AD-associated drugs from the DrugBank were mainly studied on populations in the Northern Hemisphere, and those drugs also received a higher rating by the LLM. This may indicate that the output of the LLM is not applicable to populations to the same degree. Possibly, region-specific influences on the effects of the drugs (such as climate, nutrition, ethnicity, and healthcare system) have a biased representation in the drug repurposing pipeline and need to be considered when interpreting and applying the results.

There were only clinical trials on 2,918 drugs retrievable from clinicaltrials.gov, possibly because there might not have been trials on these drugs. However, it is also possible that for those drugs that do not have a generic name but only an IUPAC classification, the data was not retrievable for technical reasons. The retrieval process and its technical limitations are extensively discussed in the code documentation and the original code. The code that was used for accessing the data is entirely available in the GitHub repository, and specific explanations are available in the ‘docs’ directory.

The AD-associated drugs from the DrugBank were mainly studied in populations in the northern hemisphere, and those drugs also received a higher rating from the LLM. As the model was trained on the available data on the internet, we wanted to pay attention to flag this potential bias inherent in the LLM. This may indicate that the DR rating results are not applicable to all populations to the same degree. We took this step to address the biases in the training data of the LLM when advocating for an AI-supported Drug Repurposing approach.

### 4.4 Conclusions

This advancement may move the synergies of ontologies, databases, and LLMs closer to each other. LLMs make it possible to enhance manual research work by automating tasks that require low levels of semantic understanding. The time and resources saved may be invested in new research. Therefore, no such pipeline would replace the critical thinking of the researcher. Still, novel thought experiments and retrospective studies could be made possible by taking up the low-level understanding of knowledge work. This approach should move us toward making better mistakes in the endeavor to find a cure for AD.

## Data Availability

All the code is available on a public GitHub repo (https://github.com/virtual-twin/allmzheimer).

## Acknowledgment

Computation has been performed on the High-Performance Cluster for Research and Clinic of the Berlin Institute of Health, Berlin, Germany.

PR acknowledges support by the Virtual Research Environment at the Charité Berlin – a node of EBRAINS Health Data Cloud, by EU Horizon Europe program Horizon EBRAINS2.0 (101147319), Virtual Brain Twin (101137289), EBRAINS-PREP 101079717, AISN – 101057655, EBRAIN-Health 101058516, Digital Europe TEF-Health 101100700, EU H2020 Virtual Brain Cloud 826421, Human Brain Project SGA2 785907; Human Brain Project SGA3 945539, German Research Foundation SFB 1436 (project ID 425899996); SFB 1315 (project ID 327654276); SFB 936 (project ID 178316478; SFB-TRR 295 (project ID 424778381); SPP Computational Connectomics RI 2073/6-1, RI 2073/10-2, RI 2073/9-1; DFG Clinical Research Group BECAUSE-Y 504745852, PHRASE Horizon EIC grant 101058240; Berlin Institute of Health and Foundation Charité.

